# Multi-color droplet digital PCR assay enables allele-specific quantification of heterogeneous genome editing outcomes

**DOI:** 10.64898/2026.08.20.745440

**Authors:** Yuga Yasuda, Yuichiro Miyaoka

## Abstract

Precise characterization of genome editing outcomes remains a major challenge because edited cell populations contain diverse alleles generated by homology-directed repair, non-homologous end joining (NHEJ), or base editing. While next-generation sequencing enables comprehensive analysis, its routine use is constrained by cost and turnaround time. Here, we developed a multi-color droplet digital PCR (ddPCR) assay that exploits six-color fluorescence detection to quantitatively distinguish multiple edited alleles within a single reaction. Using CRISPR-Cas9 and base editing model systems, we designed sequence-specific probe sets that distinguished recurrent NHEJ alleles generated by CRISPR-Cas9 editing as well as target and bystander alleles generated by base editing. The assay quantitatively resolved individual editing outcomes that could not be distinguished by conventional Sanger sequencing. Together, these results establish multi-color ddPCR as a rapid, scalable, and sequence-specific approach for quantification of genome editing outcomes across multiple editing modalities.

**Graphical abstract:** 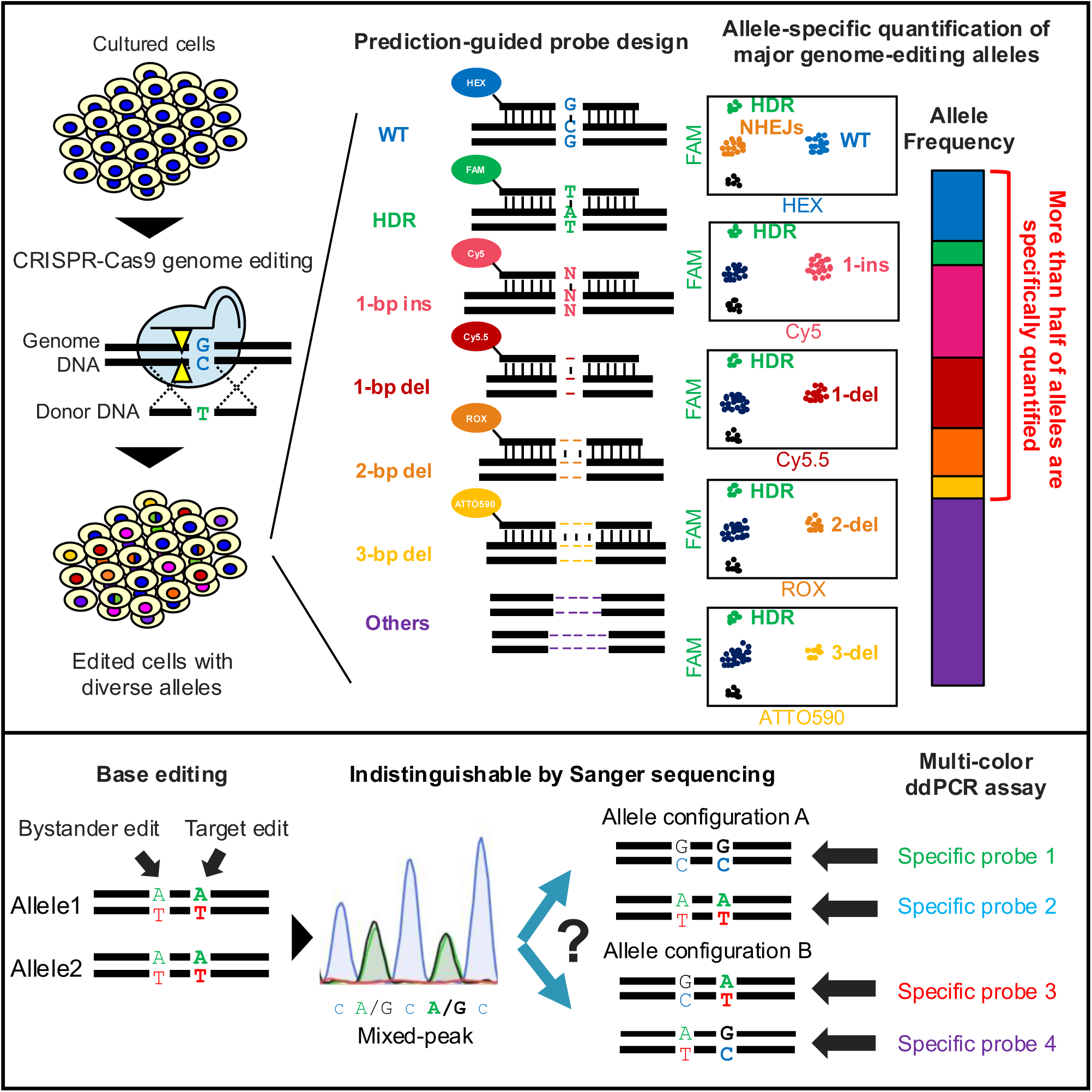

**Motivation:** Genome editing generates complex outcomes of desired edits, wild-type alleles, and heterogeneous undesired edits. Although next-generation sequencing (NGS) provides comprehensive characterization of these outcomes, routine analysis remains limited by cost, turnaround time, and analytical complexity. Existing droplet digital PCR (ddPCR) assays offer a rapid and quantitative alternative but have been unable to distinguish individual NHEJ-derived indel alleles because of the limited number of fluorescence channels available in conventional platforms. In addition, characterization of base editing outcomes is complicated by bystander edits that cannot be resolved by standard Sanger sequencing. We sought to develop a versatile multi-color ddPCR strategy capable of directly detecting and quantifying multiple editing outcomes in a single reaction, thereby providing a rapid, cost-effective alternative to sequencing-based approaches for both CRISPR-Cas9 and base editing applications.

**Highlights:**

- Six-color ddPCR assays simultaneously quantify WT, HDR, and multiple sequence-defined NHEJ alleles in a single reaction.
- The limits of detection for individual genome editing outcomes are as low as 0.08–0.30%.
- Multi-color ddPCR assays combined with machine learning–based indel prediction can capture more than 60% of editing outcomes without prior sequencing.
- Multi-color ddPCR assays enable discrimination of target and bystander modifications by base editing that cannot be resolved by Sanger sequencing.

## Introduction

CRISPR-Cas9 genome editing enables efficient and flexible genetic modification by inducing targeted double-strand breaks (DSBs) in genomic DNA. Following DSB induction, DNA repair by homology-directed repair (HDR) can introduce precise sequence modifications using a donor template with intended changes, whereas DNA repair by non-homologous end joining (NHEJ) frequently generates variable insertions and deletions (indels). As a result, genome-edited cell populations contain a complex mixture of precisely edited alleles, unedited alleles, and heterogeneous indel alleles.

Therefore, accurate and quantitative characterization of complex genome editing outcomes is essential for both basic research and translational applications. Droplet digital PCR (ddPCR) provides a sensitive and scalable approach for allele quantification without the need for extensive sequencing. Since we first utilized ddPCR for detection of genome editing outcomes^1^, several groups have also demonstrated that digital PCR can be used for sensitive quantification of genome editing outcomes^2^. In particular, our improved ddPCR-based approaches have enabled quantitative detection of edited alleles, HDR events, and NHEJ outcomes^3,4^. However, due to the two-color detection limit of commonly used ddPCR platforms, diverse indel alleles were operationally grouped as NHEJ, without being distinguished.

With recent technical advancements, six-color ddPCR platforms are now available, providing an opportunity to precisely detect multiple indel alleles. However, whether multi-color ddPCR can be used to resolve individual editing outcomes generated by genome editing has not been systematically evaluated. In parallel, machine learning–based tools such as FORECasT^5^ and inDelphi^6^ have enabled *in silico* prediction of indel patterns generated by CRISPR- Cas9. These tools accurately predict recurrent repair outcomes and may facilitate assay design without prior sequencing data. Together, these advances raise the possibility of selectively targeting and quantifying defined, high-frequency indel alleles, which would have been categorized as an NHEJ group.

Beyond DSB-based genome editing, base editing has been established as an important alternative strategy that enables precise nucleotide conversions without introducing DSBs and typically generates fewer NHEJ-derived indels^7–9^. Despite this advantage, base editing presents yet another analytical challenge. Because base editors operate within a defined editing window, unintended modifications at neighboring positions, referred to as bystander editing, frequently occur^10,11^. Moreover, adenine base editors (ABEs) have been reported to induce low-frequency cytosine-to-thymine conversions at nearby sites, generating sequence variants beyond the intended A-to-G edits^12^.

When multiple bases within an editing window are edited, or when unexpected nucleotide conversions occur, conventional Sanger sequencing cannot resolve the resulting allele configurations due to overlapping chromatogram signals^13^. Although computational methods can estimate editing frequencies from mixed sequencing traces, they do not directly resolve individual allele configurations^13^. Next-generation sequencing (NGS), although accurate, remains costly and time-consuming for routine analysis^14^. These considerations suggest that a multi-color ddPCR assay could enable quantitative analysis of individual alleles generated by base editing.

Therefore, in this study, we sought to develop a versatile, multi-color ddPCR assay capable of resolving and quantifying multiple edited alleles generated by both CRISPR-Cas9 and base editing. By integrating multiple sequence-specific probes within a single ddPCR, this approach enables deconvolution of heterogeneous allele populations and provides a scalable alternative to sequencing-based methods.

## Results

### Design of a multi-color ddPCR assay for detection of multiple indel alleles

Previous ddPCR-based quantification of CRISPR–Cas9 genome editing outcomes classified alleles into three categories—homology-directed repair (HDR), unedited or wild-type (WT), and non-homologous end joining (NHEJ). Due to the two-color detection limit of earlier ddPCR platforms, NHEJ-derived indel alleles were operationally defined as a population that was neither WT nor HDR, resulting in an aggregation of diverse indel alleles (Figure 1a). To resolve this heterogeneity, we leveraged the six-color detection capability of the QX-600 ddPCR system to distinguish individual recurrent indel alleles.

**Figure 1.**
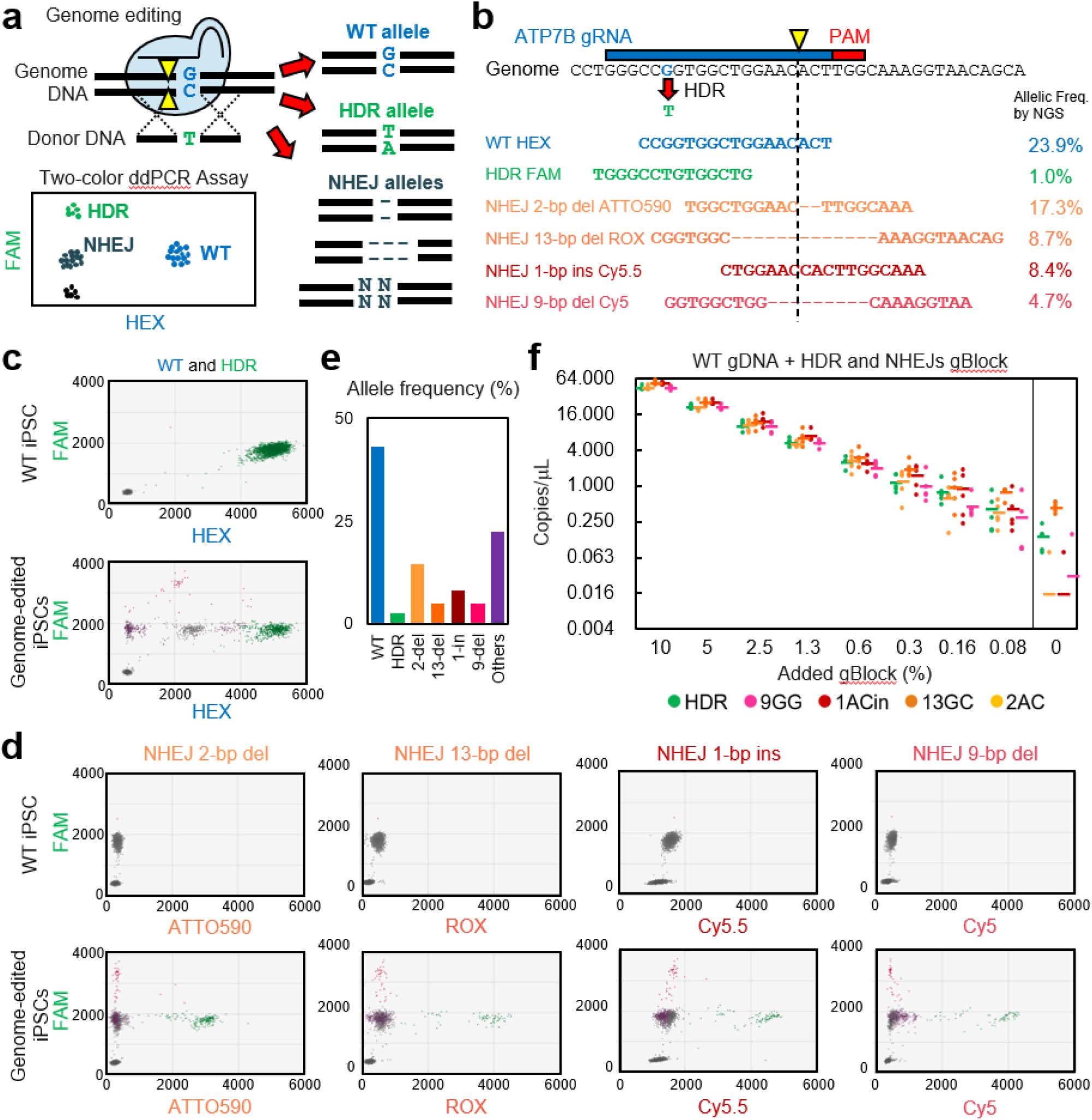
Design and validation of a multi-color ddPCR assay for detection of multiple genome editing outcomes. (a) Limitation of conventional two-color ddPCR strategies for analyzing CRISPR-Cas9 genome editing outcomes. In conventional ddPCR assays, genome editing outcomes are classified into WT, HDR, and NHEJ categories. Diverse NHEJ-derived indel alleles are grouped into a single NHEJ population. (b) Design of the ATP7B R778L multi-color ddPCR assay. A reference FAM probe, WT-specific HEX probe, HDR-specific FAM probe, and four NHEJ allele-specific probes labeled with ATTO590, ROX, Cy5.5, and Cy5 were designed to competitively hybridize to the target region. Previously determined allele frequencies by NGS are indicated. (c) Representative two-dimensional fluorescence plots of WT control genomic DNA (top) and ATP7B-edited induced pluripotent stem (iPS) cell genomic DNA (bottom) displayed on the FAM–HEX axes. WT genomic DNA exhibited a single WT population, whereas edited samples contained additional populations corresponding to HDR and NHEJ alleles. (d) Representative two-dimensional fluorescence plots showing detection of HDR and four individual NHEJ alleles in WT control genomic DNA (top) and ATP7B-edited iPS cell genomic DNA (bottom). (e) Quantification of ATP7B genome editing outcomes by multi-color ddPCR. Frequencies of WT, HDR, four NHEJ alleles, and Others populations are shown. Others population represents droplets positive only for the reference FAM probe and corresponds to edited alleles not specifically recognized by the designed allele-specific probes. (f) Quantitative performance of the multi-color ddPCR assay assessed using serial dilutions of synthetic HDR and NHEJ templates spiked into WT genomic DNA. Measured concentrations (copies/μL) are plotted against expected template fractions. The assay demonstrated linear quantification across a broad dynamic range. Limits of detection were determined by comparison with WT-only controls and are summarized in Table S1. WT, wild type; HDR, homology-directed repair; NHEJ, non-homologous end joining; ddPCR, droplet digital PCR.

We applied this strategy to the ATP7B R778L point mutagenesis as an initial test case. We had previously characterized genome editing outcomes of this point mutagenesis in human cultured cell lines^15, 16^ and induced pluripotent stem (iPS) cells^1, 3, 17, 18^ by ddPCR.

Based on these results, we designed four probes with different fluorophores targeting the most frequently observed NHEJ alleles (2-bp deletion, 13-bp deletion, 1-bp insertion, and 9-bp deletion) for the multi-color ddPCR assay. In addition to these four NHEJ-specific probes, we incorporated previously reported primers, a reference probe to detect total genomic copy number, WT-specific and HDR-specific probes in the assay^16^. We designed these probes to overlap, so that they compete with each other for hybridization to the targeted ATP7B sequence (Figure 1b).

### Multi-color ddPCR assay enables simultaneous quantification of WT, HDR, and multiple NHEJ alleles

To validate probe specificity, we generated positive-control plasmids containing the WT allele, HDR allele, and the four representative NHEJ alleles. Analysis of each plasmid with the multi-color ddPCR assay produced the expected fluorescence populations: WT plasmids generated a FAM+/HEX+ population, HDR plasmids generated a FAM++ population, 9-del plasmids generated a FAM+/Cy5+ population, 1-ins plasmids generated a FAM+/Cy5.5+ population, 13-del plasmids generated a FAM+/ROX+ population, and 2-del plasmids generated a FAM+/ATTO590+ population, respectively (Figure 1b and S1). Minimal cross-reactivity was observed among probes, demonstrating that the assay can distinguish WT, HDR, and NHEJ alleles with high specificity.

Genomic DNA extracted from iPS cells after the ATP7B R778L point mutagenesis by CRISPR-Cas9 was analyzed using the designed multi-color ddPCR assay. Control non-edited genomic DNA exhibited a single WT population positive for both FAM and HEX fluorescence (Figure 1c). In contrast, the edited samples displayed distinct populations corresponding to the induced HDR and NHEJ alleles (Figure 1d). The allele frequencies of the four NHEJ alleles, the WT allele, and the HDR allele were quantified by this multi-color ddPCR assay (Figure 1e). The overall distribution of editing outcomes was consistent with that determined previously by NGS. (Figure 1b, e).

Droplets positive only for the reference FAM probe were classified as “Others”, representing edited alleles not specifically detected by the NHEJ probes. The Others fraction accounted for 22.2% of all alleles, indicating that the assay captured nearly 80% of genome editing outcomes while simultaneously resolving WT, HDR, and major NHEJ alleles. These results demonstrate that our multi-color ddPCR assay detect multiple specific NHEJ alleles in addition to the WT and HDR alleles.

We further assessed the quantitative performance of the multi-color ddPCR assay using synthetic DNA templates containing HDR and representative NHEJ alleles (Table S1). Analysis of WT genomic DNA spiked with defined amounts of synthetic templates demonstrated linear quantification across a broad dynamic range (Figure 1f). Limits of detection (LoD) were estimated based on non-overlap of 95% confidence intervals between dilution samples and WT-only controls, yielding LoDs of 0.16% for HDR and 9-del, 0.3% for 1-ins and 13-del, and 0.08% for 2-del (Table S2).

### Multi-color ddPCR can be adapted to different genome editing targets

To confirm that the multi-color ddPCR assay can be designed flexibly for other genome editing, we tested another primer-probe set for RBM20 R636S point mutagenesis, which had also been analyzed before^3, 18^. Unlike the ATP7B R778L point mutagenesis, the RBM20 R636S point mutagenesis site was 12 bp away from the DNA cleavage site by Cas9. Therefore, we incorporated a DARK probe in addition to four sequence-specific NHEJ probes (8-bp deletion-1, 1-bp deletion, 9-bp deletion, and 8-bp deletion-2), which were determined by our previous analyses^18^ (Figure 2a). We induced the RBM20 R636S point mutation in human iPS cells and analyzed the genome editing outcomes in them by this multi-color ddPCR assay. As a result, we were able to detect induction of individual NHEJ alleles in addition to WT and HDR alleles (Figure 2b, c and S2). Quantification of the frequencies of WT, HDR, and the four induced NHEJ alleles revealed that the frequency of Others alleles accounted for 42.6% of all alleles, indicating that the assay captured nearly 60% of genome editing outcomes while simultaneously resolving WT, HDR, and major recurrent NHEJ alleles (Figure 2d). The allele frequencies determined by multi-color ddPCR assay well matched those determined by NGS previously (Figure 2a, d).

**Figure 2.**
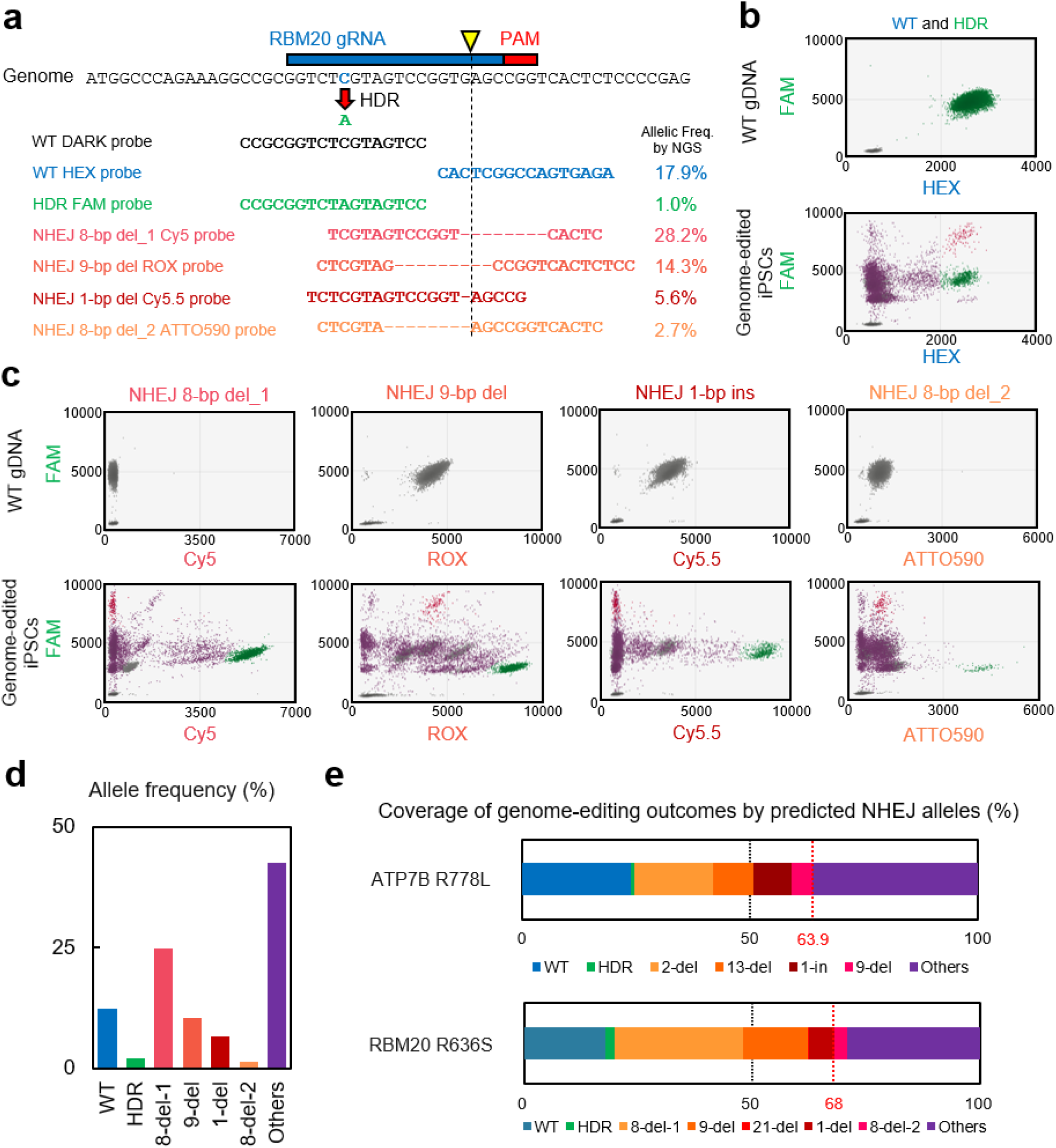
Flexible design of the multi-color ddPCR assay and prediction-guided identification of recurrent NHEJ alleles. (a) Design of the RBM20 R636S multi-color ddPCR assay. Because the target nucleotide is located 12 bp away from the Cas9 cleavage site, a DARK probe was incorporated together with a reference probe, WT-specific probe, HDR-specific probe, and four NHEJ allele-specific probes targeting the most frequently observed NHEJ alleles (8-bp deletion-1, 1-bp deletion, 9-bp deletion, and 8-bp deletion-2). Previously determined allele frequencies by NGS are indicated. (b) Representative two-dimensional fluorescence plots of WT control genomic DNA (top) and RBM20-edited induced pluripotent stem (iPS) cell genomic DNA (bottom) displayed on the FAM–HEX axes. WT genomic DNA exhibited a single WT population, whereas edited samples contained additional fluorescence populations corresponding to HDR and NHEJ alleles. (c) Representative two-dimensional fluorescence plots showing detection of four individual NHEJ alleles in WT control genomic DNA (top) and RBM20-edited iPS cell genomic DNA (bottom). (d) Quantification of RBM20 genome editing outcomes by multi-color ddPCR. Frequencies of WT, HDR, four NHEJ alleles, and Others populations are shown. Others population represents droplets positive only for the reference FAM probe and corresponds to edited alleles not specifically recognized by the sequence-specific probes. (e) Predicted coverage of genome editing outcomes by four NHEJ alleles selected using the FORECasT. The four highest-frequency NHEJ alleles predicted by FORECasT were assumed to be targets for the multi-color ddPCR assay. Together with the WT and HDR alleles, these predicted NHEJ alleles accounted for 63.9% and 67.9% of all genome editing outcomes observed for ATP7B R778L and RBM20 R636S editing, respectively, demonstrating that more than half of editing outcomes can be captured using only four predicted high-frequency NHEJ alleles.

These results demonstrated that the multi-color ddPCR can be flexibly designed for various genome editing targets.

### More than half of genome editing outcomes can be captured by the multi-color ddPCR and *in silico* prediction

Since the QX600 ddPCR system can detect up to six fluorophores simultaneously, our multi-color ddPCR assay can theoretically detect a maximum of four specific NHEJ alleles in addition to WT and HDR alleles. We therefore evaluated whether *in silico* prediction tools could be used to identify high-frequency NHEJ alleles for inclusion in a multi-color ddPCR assay in the absence of prior knowledge of editing outcomes. Genome editing outcomes for ATP7B and RBM20 were predicted using FORECasT^5^ and inDelphi^6^ and compared with NGS data obtained previously in our laboratory (Table S3).

For ATP7B R778L editing, the top four NHEJ alleles predicted by both FORECasT and inDelphi corresponded to the four most frequently observed NHEJ alleles identified by NGS. Together with the WT and HDR alleles, these four NHEJ alleles accounted for 63.9% of all observed editing outcomes (Figure 2e).

For RBM20 R636S editing, three of the top four alleles predicted by FORECasT (ranked first, second, and fourth) were among the four most abundant NHEJ alleles observed by NGS. In contrast, the third-ranked FORECasT prediction corresponded to the 74th most abundant allele in the NGS dataset (allele frequency, 0.1%), whereas the fourth most abundant NGS allele was ranked 27th by FORECasT (Table S3). Nevertheless, the four highest-frequency NHEJ alleles predicted by FORECasT, together with the WT and HDR alleles, accounted for 68% of all observed editing outcomes (Figure 2e). These results suggest that a multi-color ddPCR assay incorporating the high-frequency NHEJ alleles predicted *in silico* can capture more than half of genome editing outcomes without prior comprehensive sequencing analysis.

### Multi-color ddPCR assay distinguishes target, bystander, and unexpected base editing outcomes

Next, we applied the multi-color ddPCR assay to analyze base editing outcomes. When both target and bystander bases are edited, multiple allele configurations can be generated in a single cell. Sanger sequencing cannot distinguish between different combinations of base editing outcomes, as overlapping chromatogram signals obscure allele-specific patterns (Figure 3a). We therefore evaluated whether the multi-color ddPCR assay could distinguish individual base editing outcomes.

**Figure 3.**
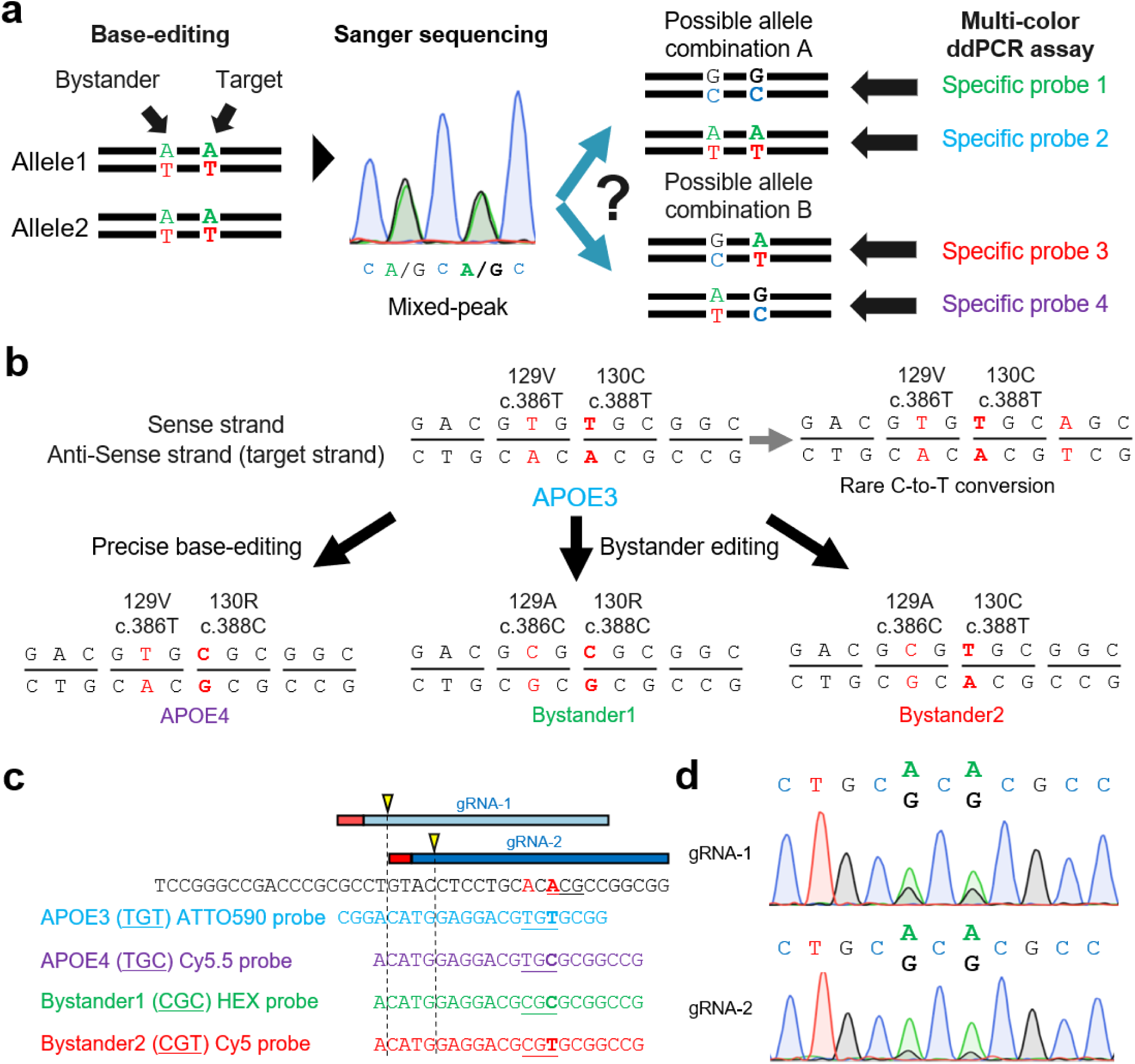
Design of a multi-color ddPCR assay for analysis of base editing outcomes. (a) Limitations of Sanger sequencing and the advantages of multi-color ddPCR analysis of base editing outcomes. Distinct allele configurations generated by target and bystander editing can produce similar mixed chromatogram signals that cannot be uniquely resolved by Sanger sequencing. In contrast, multi-color ddPCR using sequence-specific probes enables discrimination of individual allele configurations. (b) Predicted editing outcomes generated during ABE8e-mediated conversion of APOE3 to APOE4. The target adenine corresponding to c.388 and a nearby adenine corresponding to c.386 are located within the same editing window. Editing of c.388 generates the APOE4 allele, whereas editing of c.386 generates bystander alleles. Rare cytosine-to-thymine conversion events previously reported for adenine base editors might generate additional unexpected editing products. (c) Design of the APOE multi-color ddPCR assay. Two guide RNAs (gRNA-1 and gRNA-2) were used for ABE8e-mediated editing. Sequence-specific probes were designed to detect APOE3 (WT), APOE4, Bystander-1, and Bystander-2 alleles within a single PCR amplicon. (d) Representative Sanger sequencing chromatograms of genomic DNA from cells edited using gRNA-1 and gRNA-2. Mixed peaks were observed at both the target and bystander positions, confirming successful editing but preventing unambiguous determination of individual allele configurations.

As a test case, we selected the APOE locus, in which the APOE3 (c.388T) and APOE4 (c.388C) alleles are distinguished by the rs429358 variant (c.388T>C). To generate the APOE4 allele from APOE3 using ABE8e^12^, we targeted the adenine on the complementary strand corresponding to c.388. A second adenine corresponding to c.386 was located within the editing window. Editing of this position generates a bystander edit, resulting in multiple possible allele configurations (Figure 3b). We performed adenine base editing from APOE3 to APOE4 using ABE8e and two distinct guide RNAs (gRNA-1 and gRNA-2). We designed sequence-specific fluorescent probes targeting the expected alleles generated by base editing, including the intended APOE4 allele and bystander-edited alleles (Figure 3c). A reference FAM probe was included to quantify the total copy number of the target locus, and all probes were incorporated within a single PCR amplicon.

Sanger sequencing confirmed successful base editing with both gRNAs, revealing that gRNA-1 exhibited slightly higher overall activity than gRNA-2 and that both the target APOE4 site and the bystander site were edited with comparable efficiencies for each gRNA (Figure 3d). However, individual allele configurations could not be resolved by Sanger sequencing.

In contrast, the multi-color ddPCR assay enabled clear separation of distinct base-edited allele populations. In the wild-type control sample, only an ATTO590-positive WT population was detected. In the genomic DNA extracted from base-edited cells, additional HEX-, Cy5-, and Cy5.5-positive droplet populations were observed at distinct positions not present in the control sample, corresponding to bystander-edited and APOE4 alleles (Figure 4a and S3). For gRNA-1, the HEX-positive Bystander-1 allele was detected at 24.8%, whereas the Cy5-positive Bystander-2 allele and the Cy5.5-positive APOE4 allele were detected at frequencies below 0.2%. In contrast, gRNA-2 editing resulted in 9.9% HEX-positive Bystander-1 alleles, 5.3% Cy5-positive Bystander-2 alleles, and 4.6% Cy5.5-positive APOE4 alleles (Figure 4b).

**Figure 4.**
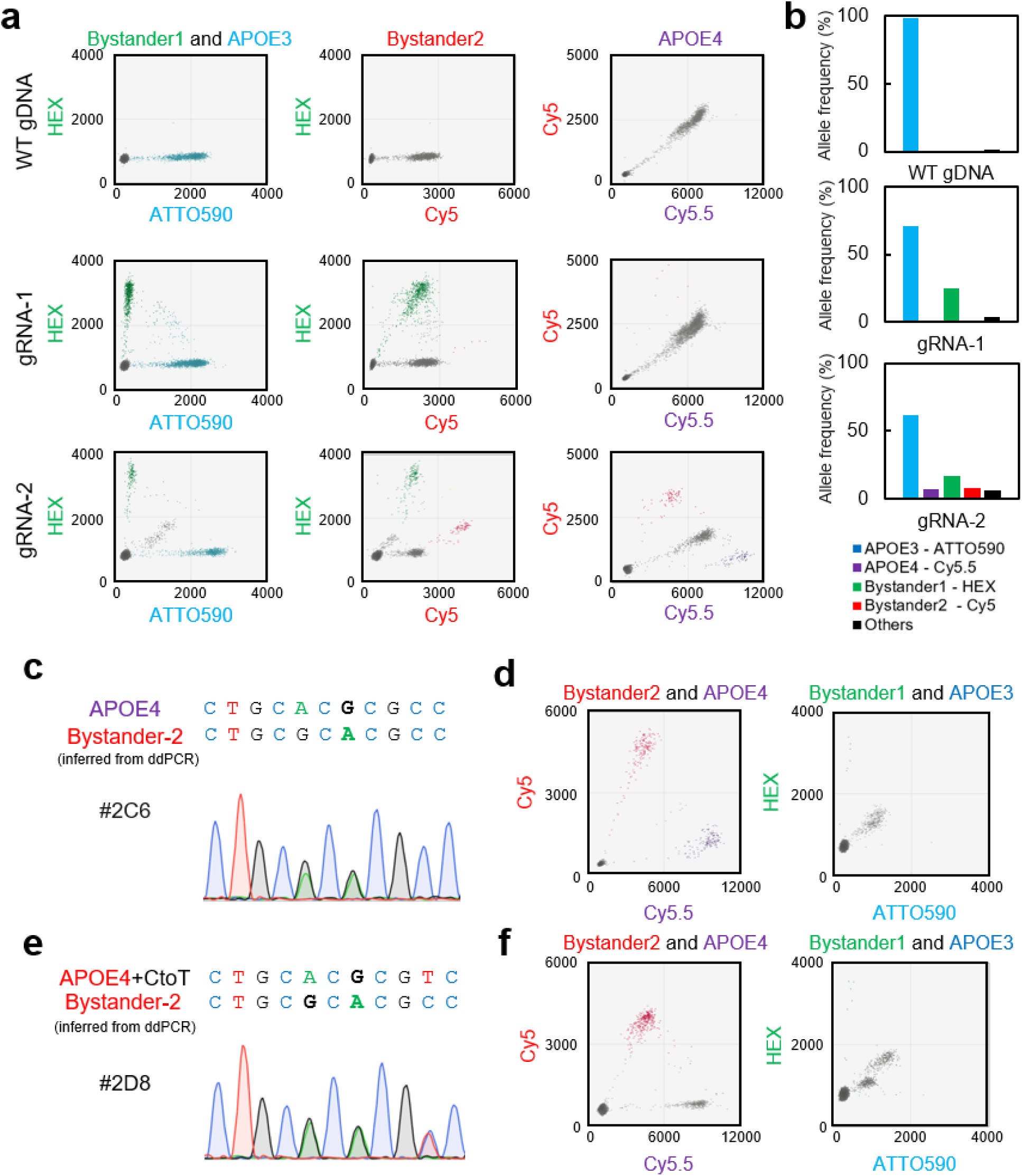
Multi-color ddPCR enables quantitative analysis of base editing outcomes at population and clonal levels. (a) Representative two-dimensional fluorescence plots of WT APOE3 genomic DNA and genomic DNA from cells edited with ABE8e using gRNA-1 or gRNA-2. Distinct fluorescence populations corresponding to APOE4, Bystander-1, and Bystander-2 alleles were detected in edited samples but not in the WT control. Fluorescence amplitudes of ATTO590, HEX, Cy5, and Cy5.5 channels are shown. (b) Quantification of base editing outcomes by multi-color ddPCR. Frequencies of APOE3 (WT), APOE4, Bystander-1, Bystander-2, and Others population were determined for WT genomic DNA and cells edited using gRNA-1 or gRNA-2. (c) Sanger sequencing chromatogram of clone #2C6. The allele configurations shown above the chromatogram were inferred from ddPCR-based genotyping. (d) Two-dimensional fluorescence plots of clone #2C6. Distinct populations corresponding to APOE4 and Bystander-2 alleles were detected, whereas Bystander-1 and APOE3 populations were absent. (e) Sanger sequencing chromatogram of clone #2D8. The allele configurations shown above the chromatogram were inferred from ddPCR-based genotyping. (f) Two-dimensional fluorescence plots of clone #2D8. A population corresponding to the APOE4+C-to-T allele was detected together with a Bystander-2 population. The APOE4+C-to-T population exhibited reduced Cy5.5 fluorescence intensity compared with the APOE4 population detected in clone #2C6, consistent with reduced hybridization of the APOE4-specific probe caused by the additional nucleotide substitution.

To validate allele assignments, we isolated individual clones and analyzed them by multi-color ddPCR. Clone #2C6 was identified as a heterozygote carrying APOE4 and Bystander-2 alleles, a configuration that could not be distinguished by Sanger sequencing (Figure 4c, d). In addition, clone #2D8 was identified as a heterozygote carrying an APOE4+C-to-T allele in combination with a Bystander-2 allele (Figure 4e, f). Although adenine base editors are designed to mediate A-to-G conversions, low-frequency cytosine-to-thymine editing has been reported previously^12^. Notably, the APOE4+C-to-T population exhibited reduced Cy5.5 fluorescence intensity compared with the APOE4 population detected in clone #2C6, consistent with decreased hybridization of the APOE4-specific probe caused by the additional nucleotide substitution. Detection of this unexpected allele configuration further illustrates the ability of the multi-color ddPCR assay to resolve complex editing outcomes beyond the anticipated target and bystander edits.

Together, these results show that the multi-color ddPCR assay can detect and quantify base editing outcomes at both population and clonal levels, including predicted bystander edits and unexpected editing products generated within the editing window.

## Discussion

Quantification of genome editing outcomes has traditionally relied mainly on sequencing-based approaches, including Sanger sequencing^13^ and NGS^14^. Sanger sequencing is limited in its sensitivity and quantitative performance, and NGS is associated with higher cost and longer turnaround times. We have shown that ddPCR offers a scalable and quantitative alternative for targeted analysis of genome editing outcomes, but earlier implementations were constrained by a limited number of available fluorophores, preventing resolution of heterogeneous allele populations such as diverse NHEJ-derived indels^1–4^.

In this study, we demonstrate that integration of multi-color fluorescence detection and sequence-specific probe design substantially expands the analytical capability of ddPCR. For CRISPR–Cas9–mediated genome editing, this approach enables quantitative detection of four specific indel alleles that were previously aggregated into a single NHEJ group. In addition, we show that prediction-guided selection of high-frequency edited alleles using machine learning tools such as FORECasT and inDelphi can facilitate assay development, and the resulting multi-color ddPCR assays can quantify more than half of genome editing outcomes.

Importantly, we further demonstrate that the same multi-color ddPCR framework can be extended to base editing. By targeting defined sequence configurations generated within the editing window, this approach enables discrimination and quantification of targeted and bystander alleles that cannot be resolved by Sanger sequencing. This capability provides practical advantages for evaluating guide RNA performance and interpreting base editor specificity. Notably, the assay also resolved an APOE4 allele carrying an additional C-to-T substitution, consistent with previous reports that adenine base editors can induce low-frequency cytosine editing. This observation highlights the utility of multi-color ddPCR for identifying complex and unanticipated editing outcomes.

The utility of this approach is expected to be greatest in applications that require repeated evaluation of editing outcomes at a predefined genomic locus, such as guide RNA optimization, donor-template design, clonal characterization of genome-edited cells, and optimization of genome editing conditions for gene therapy development. In such settings, the same probe set can be reused across large numbers of samples, providing a scalable alternative to sequencing-based approaches. Conversely, because the assay relies on locus-specific probe design, NGS remains preferable for discovery-oriented studies involving diverse genomic targets or previously uncharacterized editing outcomes.

Together, these findings establish our multi-color ddPCR assay for rapid, quantitative, and sequence-specific analysis of genome editing outcomes across multiple editing modalities. This multi-color ddPCR assay complements conventional sequencing-based methods by enabling targeted and scalable quantification of predicted or predefined alleles, thereby enhancing genome editing experiments.

## Limitations of the study

The number of alleles that can be simultaneously resolved is constrained by the number of available fluorescence channels and probe design. Moreover, rare or unanticipated editing outcomes not represented by the probe set will not be detected, positioning this assay as a targeted rather than discovery-based method. For applications requiring comprehensive discovery of editing outcomes or analysis of previously uncharacterized loci, sequencing-based approaches remain preferable.

## STAR * Methods

### Key resources table

| REAGENT OR RESOURCE | SOURCE | IDENTIFIER |
| --- | --- | --- |
| Chemicals, peptides, and recombinant proteins |  |  |
| Growth factor-reduced Matrigel | Corning | 356231 |
| mTeSR Plus | STEMCELL Technologies | ST-100-0276 |
| Accutase | Nacalai Tesque | 12679-54 |
| Y-27632 | Fujifilm Wako Pure Chemical | 18188-04 |
| Lipofectamine Stem Reagent | Thermo Fisher Scientific | STEM00008 |
| Opti-MEM | Thermo Fisher Scientific | 10829018 |
| Puromycin | Nacalai Tesque | 29455-54 |
| Blasticidin | Wako | 029-18701 |
| FastDigest HindIII | Thermo Fisher Scientific | FD0505 |
| Critical commercial assays |  |  |
| ddPCR Multiplex Supermix | Bio-Rad | 12005909 |
| FastGene Gel/PCR Extraction Kit | NIPPON Genetics | FG-91302 |
| pGEM-T Easy Vector System | Promega | A1360 |
| Experimental models: Cell lines |  |  |
| WTC11 human iPS line | Coriell Institute | GM25256 |
| HPS0076 human iPS line | RIKEN BRC | HPS0076 |
| Oligonucleotides |  |  |
| ATP7B R778L ddPCR primer/probe set | This study | Table S5 |
| RBM20 R636S ddPCR primer/probe set | This study | Table S5 |
| APOE ddPCR primer/probe set | This study | Table S5 |
| ATP7B gRNA | Takahashi et al. 2025<sup>15</sup> | Table S4 |
| RBM20 gRNA | Takahashi et al. 2025<sup>15</sup> | Table S4 |
| APOE gRNA-1 | This study | Table S4 |
| APOE gRNA-2 | This study | Table S4 |
| Recombinant DNA |  |  |
| px459-HypaCas9 | Kato-Inui et al. 2018<sup>16</sup> | RRID: Addgene_108294 |
| NG-ABE8e | Richter et al. 2020<sup>19</sup> | RRID: Addgene_138491 |
| pGB | Kato-Inui et al. 2024<sup>20</sup> | RRID: Addgene_204744 |
| ATP7B WT control plasmid | This study | N/A |
| ATP7B HDR control plasmid | This study | N/A |
| ATP7B representative NHEJ control plasmids | This study | N/A |
| RBM20 WT control plasmid | This study | N/A |
| RBM20 HDR control plasmid | This study | N/A |
| RBM20 representative NHEJ control plasmids | This study | N/A |
| Software and algorithms |  |  |
| QX Manager Software | Bio-Rad | https://www.bio-rad.com/en-jp/life-science/digital-pcr/qx-software |
| FORECasT | Allen et al. 2018<sup>5</sup> | https://elixir.ut.ee/forecast/ |
| inDelphi | Shen et al. 2018<sup>6</sup> | https://indelphi.giffordlab.mit.edu/ |
| Other |  |  |
| QX600 Droplet Digital PCR System | Bio-Rad | 17007769 |
| C1000 Touch Thermal Cycler | Bio-Rad | 1851148 |

### Experimental model and study participant details Human induced pluripotent stem cells

The WTC11 human iPSC line (Coriell Institute, GM25256; male) and the HPS0076 human iPSC line (RIKEN BRC; female) were used in this study. Cell culture conditions are described in Method Details. The sex of the donors from which these iPSC lines were derived was not evaluated as a biological variable because this study focused on development and validation of a genome editing assay platform.

### Quantification and statistical analysis

Droplet fluorescence data were analyzed using QX Manager Software (Bio-Rad), and allele frequencies were calculated from concentration values generated by the software based on Poisson statistics as described in Method Details.

Limits of detection (LoDs) were determined using dilution series experiments. LoDs were defined as the lowest allele frequency at which the 95% confidence intervals of dilution samples did not overlap with those of WT-only controls. Details regarding sample numbers and replicate measurements are provided in the corresponding figure legends. No formal hypothesis-testing statistical analyses were performed in this study.

## Method details

### Plasmids and Oligonucleotides

The px459-HypaCas9 plasmid, which expresses HypaCas9 and the puromycin resistance gene for genome editing, has been described previously^16^ (Addgene Plasmid #108294). The guide RNAs (gRNAs) and single-stranded oligodeoxynucleotide (ssODN) donors used for ATP7B and RBM20 genome editing were described previously^15^. To generate positive-control plasmids for the multi-color ddPCR assays, genomic DNA was isolated from previously established iPSC clones harboring the edited alleles of ATP7B or RBM20. Genomic regions encompassing the target loci were amplified by PCR and subsequently cloned into the pGEM-T Easy vector (Promega) using TA cloning according to the manufacturer’s instructions. Positive-control plasmids containing the WT allele, HDR allele, and representative NHEJ alleles were used for probe validation and assay development.

For base editing experiments, NG-ABE8e and pGB plasmids were used. NG-ABE8e was a gift from David Liu^19^ (Addgene plasmid #138491; http://n2t.net/addgene:138491; RRID: Addgene_138491), and pGB has been described previously^20^ (Addgene Plasmid #204744). All oligonucleotide and gRNA sequences used in this study are provided in Table S4.

### Culture and Transfection of iPSC

The WTC11 human iPSC line and the HPS0076 human iPSC line were used for genome editing and base editing experiments, respectively. iPS cells were cultured on growth factor–reduced Matrigel (Corning) in mTeSR Plus medium (STEMCELL Technologies) supplemented with 1% penicillin–streptomycin (Nacalai Tesque) at 37°C in a humidified atmosphere containing 5% CO₂. The medium was changed every other day, and cells were passaged at approximately 80% confluence. For passaging, cells were dissociated with Accutase (Nacalai Tesque) at 37°C for 5 minutes. When the cell density was below 10% confluence, 10 µM Y-27632 (Fujifilm Wako Pure Chemical) was added to the medium.

For transfection of plasmids, 4.0 × 10⁴ (for genome editing) or 5.0 × 10⁴ (for base editing) iPS cells were seeded into a Matrigel-coated 24-well plate. The next day, at least one hour before transfection, the medium was replaced with 500 µL of fresh mTeSR Plus with Ri. Transfections were performed using 2 μL of Lipofectamine Stem Reagent (Thermo Fisher Scientific) and 25 μL of Opti-MEM (Thermo Fisher Scientific) together with 450 ng of pX459 plasmid and 50 ng of ssODN donor (for genome editing) or 250 ng of NG-ABE8e and pGB plasmids (for base editing).

Antibiotic selection was performed using 0.5 μg/mL puromycin (Nacalai Tesque) for genome-edited iPSCs or 10 μg/mL Blasticidin (Wako) for base-edited iPSCs for 24 to 72 hours after transfection.

### Droplet digital PCR

Genomic DNA (100–180 ng) or plasmid DNA (0.025 pg) was used as a template for ddPCR analysis. Primer and probe mixtures were prepared according to Table S5. Reactions were assembled in a final volume of 24 μL containing 4× ddPCR Multiplex Supermix (Bio-Rad, #12005909), template DNA, primers and probes, and Milli-Q water. For genomic DNA samples, 0.48 μL of FastDigest HindIII (Thermo Fisher Scientific, #FD0505) was included in the reaction to digest genomic DNA before amplification. Plasmid DNA samples were analyzed without restriction enzyme digestion. Droplets were generated according to the manufacturer’s instructions using the QX600 Droplet Digital PCR System (Bio-Rad). Following droplet generation, PCR amplification was performed on a C1000 Touch Thermal Cycler (Bio-Rad).

Two thermal cycling programs were used depending on the target locus: Condition 1 (RBM20):

1. 95°C for 10 min
2. 94°C for 30 s
3. 59°C for 1 min
4. Repeat steps 2 and 3 for 39 additional cycles
5. 98°C for 10 min

Condition 2 (ATP7B and APOE):

1. 95°C for 10 min
2. 94°C for 30 s
3. 58°C for 1 min
4. 72°C for 2 min
5. Repeat steps 2–4 for 39 additional cycles
6. 98°C for 10 min

For both programs, the ramp rate was set to 2°C/s for all steps.

### Detection and quantification of target alleles by ddPCR

Droplet fluorescence data were analyzed using QX Manager Software (Bio-Rad). ATP7B and RBM20 assays were analyzed using the Direct Quantification experiment type with the Amplitude Multiplex assay setting. APOE assays were analyzed using the Direct Quantification experiment type with the Single Target per Channel assay setting.

Population gating was performed sequentially using no-template controls (NTCs), WT genomic DNA controls, and fluorescence signatures generated by allele-specific probes. First, droplets lacking detectable fluorescence signals were defined as the non-template population based on Milli-Q water samples (NTCs). All remaining droplets positive for the reference FAM probe were classified as template-containing droplets.

For ATP7B and RBM20 assays, non-edited WT genomic DNA was used to define the WT population, which was characterized by HEX fluorescence in addition to the reference FAM signal. Within QX Manager Software, the WT population was defined based solely on the WT-specific HEX signal and was gated as FAM−/HEX+. The reference FAM signal associated with this population was then excluded from subsequent gating steps. By iteratively excluding assigned probe-specific populations from the initial FAM-positive population, the remaining droplets could be further classified according to their allele-specific fluorescence signatures.

HDR alleles were identified as droplets exhibiting approximately two-fold higher FAM fluorescence intensity than the WT population, reflecting simultaneous hybridization of both the reference FAM probe and the HDR-specific FAM probe (FAM++ population).

After exclusion of the WT and HDR populations, droplets corresponding to individual NHEJ alleles were sequentially identified using the fluorescence channels that provided the greatest separation from previously assigned populations. In general, a greater number of probe–template mismatches resulted in better spatial separation of droplet populations on two-dimensional fluorescence plots.

For APOE base editing assays, the WT population was defined using non-edited APOE3 genomic DNA and the WT-specific ATTO590 signal. Populations corresponding to the designed base editing outcomes were subsequently identified according to their allele-specific fluorescence signatures using the same sequential gating strategy described above.

Droplets remaining positive only for the reference FAM probe after all sequence-specific populations had been assigned were classified as the Others population, representing alleles not recognized by any of the designed sequence-specific probes.

Following population assignment by sequential gating on two-dimensional fluorescence plots, the concentration of each population (copies/μL) was obtained from the “Concentration” data exported by QX Manager Software (Bio-Rad).

Allele frequencies were calculated by dividing the concentration of each allele-specific population by the summed concentrations of all defined populations within the same assay. For ATP7B and RBM20 genome editing assays, the total allele concentration was calculated as the sum of the WT, HDR, four NHEJ, and Others population.

For example, the frequency of the WT allele was calculated as:

Freq (WT) = Conc (WT) / [Conc (WT) + Conc (HDR) + Conc (NHEJ1) + Conc (NHEJ2) + Conc (NHEJ3) + Conc (NHEJ4) + Conc (Others)]

where Conc (WT), Conc (HDR), Conc (NHEJ1–4), and Conc (Others) represent the concentrations (copies/μL) of the corresponding populations reported by QX Manager Software.

The frequencies of HDR, NHEJ, and base editing alleles were calculated using the same approach by substituting the corresponding allele-specific population concentration into the numerator. The sum of all allele frequencies within a sample was defined as 100%.

### *In silico* prediction of NHEJ alleles

Genome editing outcomes were predicted using the web-based versions of FORECasT and inDelphi. For each target locus, a 50-bp genomic sequence surrounding the predicted Cas9 cleavage site was submitted to the prediction tools. FORECasT predictions were generated using the default settings provided by the web server. For inDelphi analysis, HEK293 was selected as the cell type because a human iPSC-specific option was not available, and all other parameters were left at their default values. Predicted indel alleles were ranked according to their predicted frequencies.

### Freezing iPSCs and gDNA extraction from iPSCs in a 96-well plate for sib-selection

Base-edited iPSC clones were isolated using a sib-selection strategy previously established for the enrichment and recovery of precisely edited human iPSCs^1^. Briefly, following transfection of iPSCs with base editing components, edited cell populations were subdivided into multiple sibling cultures and screened for the desired editing event by multi-color ddPCR assay. Cultures enriched for the target edits were subsequently subjected to additional rounds of sib selection and clonal isolation until genetically homogeneous clones were obtained.

### Sanger sequencing

Genomic DNA was isolated from iPSC clones obtained by sib selection after the APOE base editing. The APOE target region was amplified by PCR using the following primers: APOE_Genotyping_Fw (5′-AGCCCTTCTCCCCGCCTCCCACTGT-3′) and APOE_Genotyping_Rv (5′-CTCCGCCACCTGCTCCTTCACCTCG-3′). PCR products were purified using FastGene Gel/PCR Extraction Kit (NIPPON Genetics) and subjected to Sanger sequencing for genotyping analysis. Sequencing using the APOE_Genotyping_Fw primer was performed by Eurofins Genomics (Tokyo, Japan). Sequence chromatograms were analyzed to confirm the presence of the intended APOE edits and bystander editing events.

## Resource availability

### Lead contact

Further information and requests for resources and reagents should be directed to and will be fulfilled by the lead contact, Yuichiro Miyaoka.

### Materials availability

Plasmids generated in this study are available from the lead contact upon reasonable request.

### Data and code availability

All data reported in this study are available within the paper and its Supplemental Information files. Previously generated NGS datasets used in this study are described in our previous paper (Takahashi et al. 2025^18^) and available from the lead contact upon request.

This study does not report original code.

Any additional information required to reanalyze the data reported in this study is available from the lead contact upon request.

## Acknowledgments

We thank G. Takahashi, M. Maeda, and K. Shinozaki for providing genomic DNA samples derived from genome-edited iPSCs and previously generated NGS datasets used in this study. We thank Satoshi Ito and Katsunori Hironaka (Bio-Rad Laboratories) for valuable technical advice regarding ddPCR assay design, data analysis, and gating strategies. We also thank all lab members for helpful discussions and technical support.

This work was supported by the Japan Society for the Promotion of Science (JSPS) Grant-in-Aid for Challenging Research (Pioneering) (24K21954), JSPS Grant-in-Aid for Scientific Research (B) (24K02028), and the Japan Agency for Medical Research and Development (AMED) (24bm1423010h0002).

## Author contributions

Conceptualization, Y.Y. and Y.M.; Methodology, Y.Y. and Y.M.; Investigation, Y.Y.; Formal Analysis, Y.Y.; Writing – Original Draft, Y.Y.; Writing – Review & Editing, Y.Y. and Y.M.; Supervision, Y.M.

## Declaration of generative AI and AI-assisted technologies in the writing process

During the preparation of this work, the authors used generative AI tools, including Microsoft Copilot and ChatGPT, for language editing and manuscript refinement. The authors reviewed and revised all AI-assisted content and take full responsibility for the accuracy and integrity of the manuscript.

