## Supplemental Figure S1-3 and Table S1-5 for "Multi-color droplet digital PCR assay enables allele-specific quantification of heterogeneous genome editing outcomes"

The PDF file includes Figures S1-S3 and Table S1-S5.

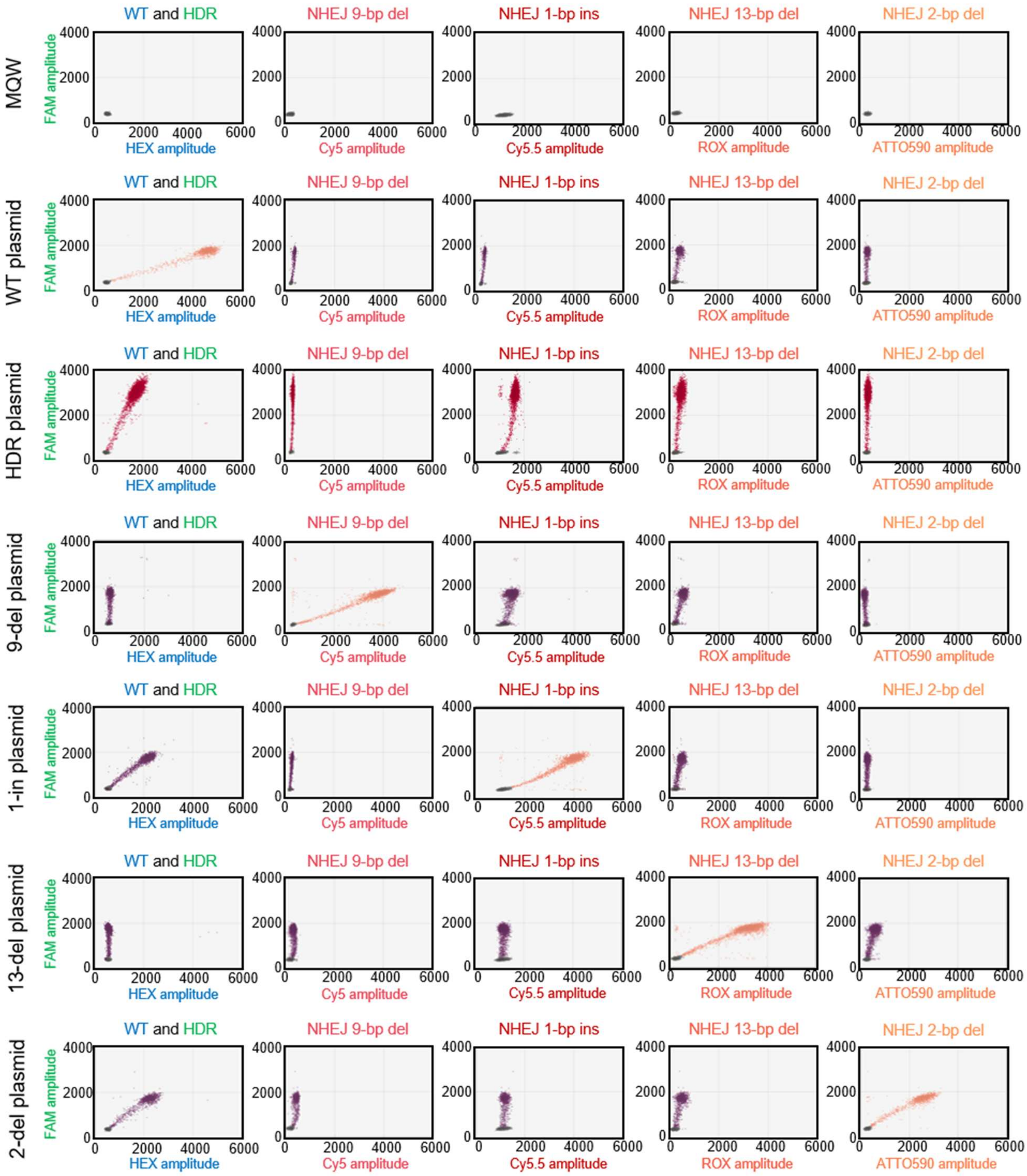


Figure S1 Validation of probe specificity for ATP7B editing using positive-control plasmids. Positive-control plasmids containing the WT allele, HDR allele, and representative NHEJ alleles (9-del, 1-ins, 13-del, and 2-del) were analyzed using the multi-color ddPCR assay. WT, HDR, 9-del, 1-ins, 13-del, and 2-del plasmids generated FAM+/HEX+, FAM++, FAM+/Cy5+, FAM+/Cy5.5+, FAM+/ROX+, and FAM+/ATTO590+ populations, respectively. Each plasmid produced the expected fluorescence cluster with no or minimal signal in non-target channels, confirming the specificity of the allele-specific probes and the ability of the assay to discriminate WT, HDR, and NHEJ alleles.

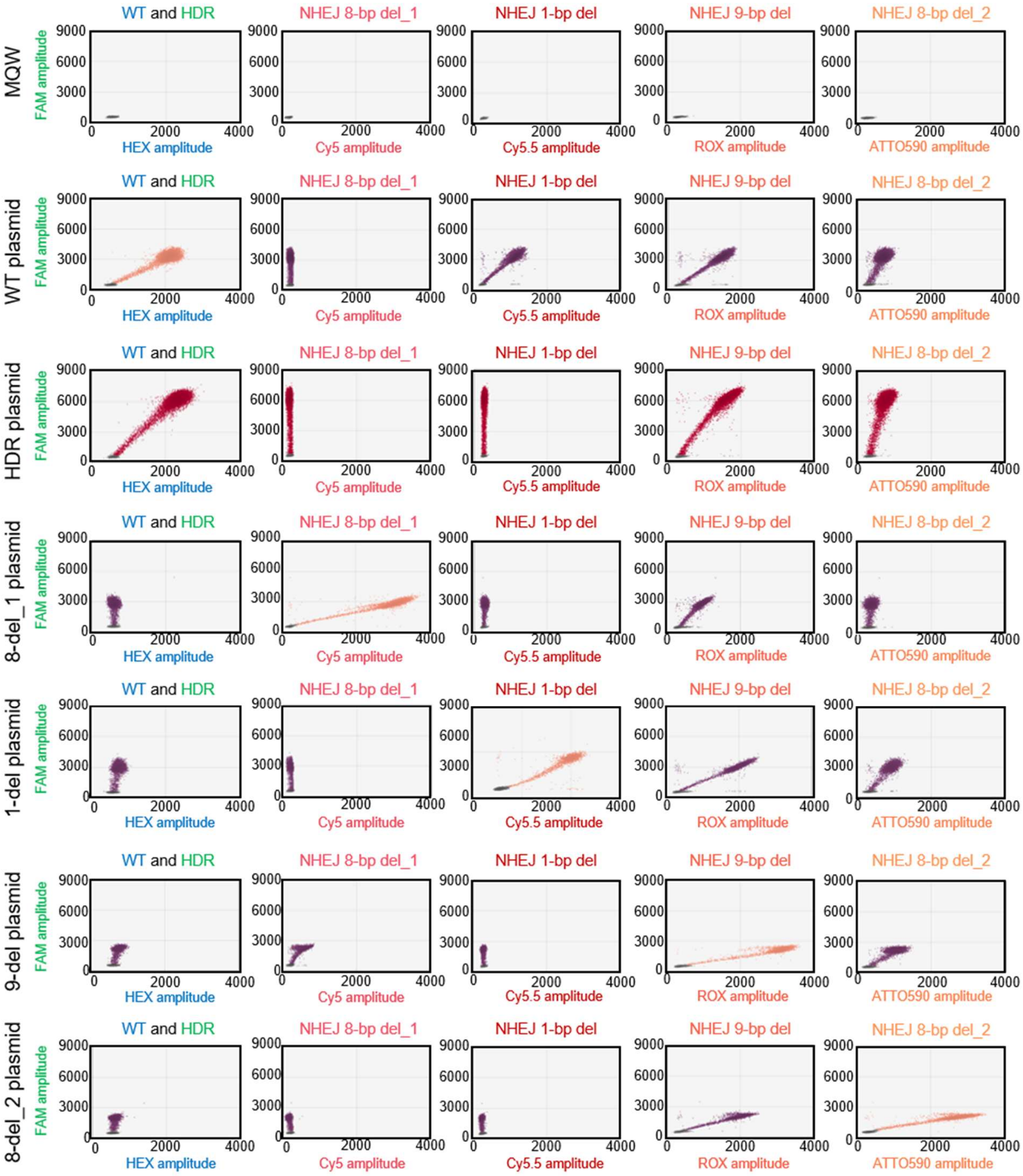


Figure S2 Validation of probe specificity for RBM20 editing using positive-control plasmids. Positive-control plasmids containing the WT allele, HDR allele, and representative NHEJ alleles (8-del_1, 1-del, 9-del, and 8-del_2) were analyzed using the multi-color ddPCR assay. WT, HDR, 8-del_1, 1-del, 9-del, and 8-del_2 plasmids generated FAM+/HEX+, FAM++, FAM+/Cy5+, FAM+/Cy5.5+, FAM+/ROX+, and FAM+/ATTO590+ populations. Each plasmid produced the expected fluorescence cluster with no or minimal signal in non-target channels, confirming the specificity of the allele-specific probes and the ability of the assay to discriminate WT, HDR, and NHEJ alleles.

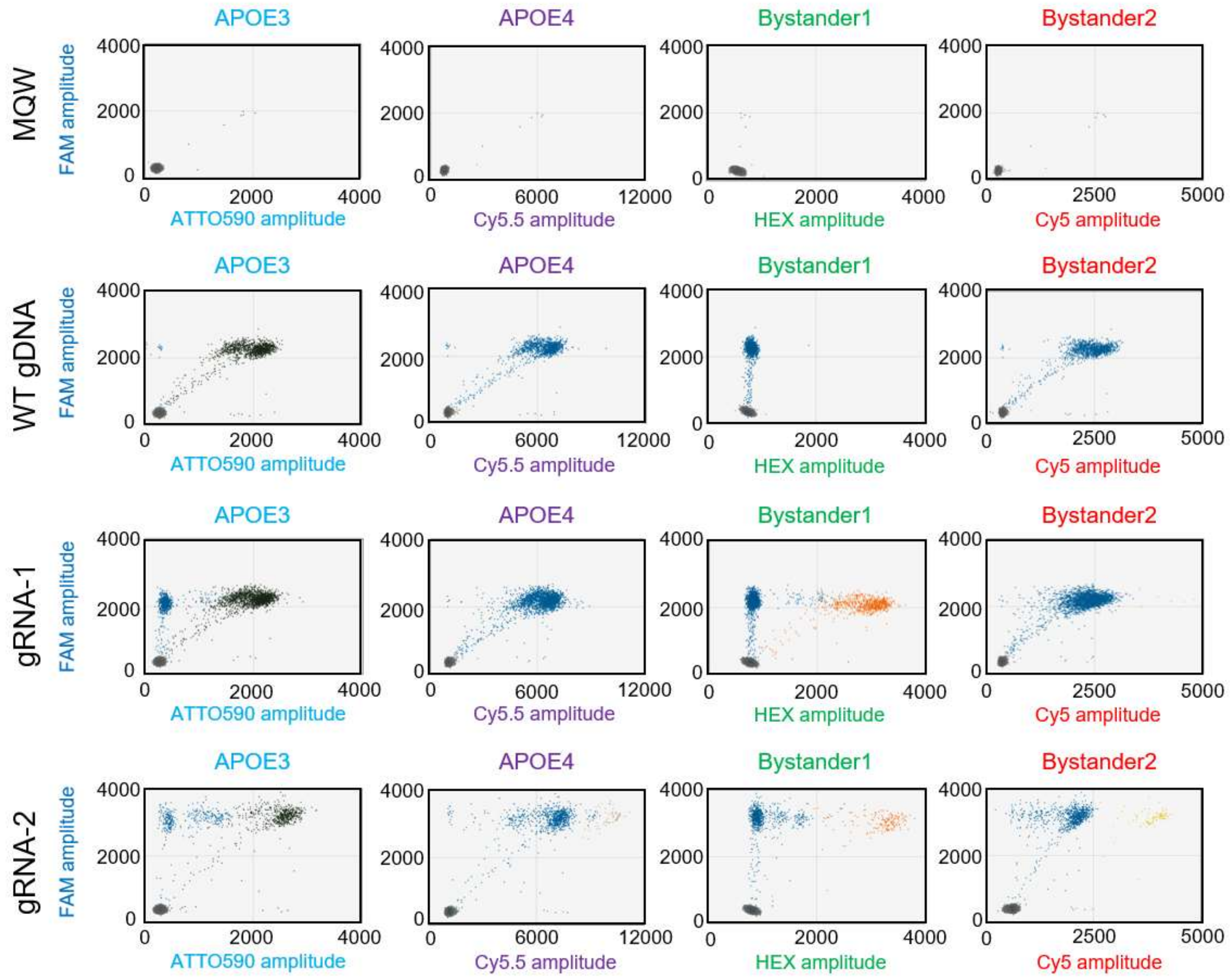


Figure S3. Reference FAM fluorescence and allele-specific fluorescence signals used for identification of APOE base-editing outcomes.

Representative two-dimensional fluorescence plots of no-template control (MQW), WT genomic DNA, and genomic DNA from cells edited with ABE8e using gRNA-1 or gRNA-2. Fluorescence amplitudes of the reference FAM probe are plotted against the allele-specific probe channels corresponding to APOE3 (ATTO590), APOE4 (Cy5.5), Bystander-1 (HEX), and Bystander-2 (Cy5).

Consistent with the competitive probe design, APOE3, APOE4, Bystander-1, and Bystander-2 populations were detected as double-positive populations for both the reference FAM probe and the corresponding allele-specific probe. No allele-specific populations were observed in no-template controls. WT genomic DNA exhibited only the APOE3 population, whereas additional APOE4 and bystander-editing populations were detected in base-edited samples. WT alleles showed minimal reactivity toward probes containing two mismatches and weak reactivity toward probes containing a single mismatch. Nevertheless, APOE4 and Bystander-2 alleles produced substantially greater fluorescence amplitudes than WT alleles, resulting in clearly separable droplet populations.

| gBlock synthetic DNA | |
|---|---|
| 9-del | CACTGTCCTTGTCTTTCAGCTCCTCGGTGGGTGGTACTTCTACGTTCAGGCCTACAAATCTC<br>TGAGACACAGGTCAGCCAACATGGACGTGCTCATCGTCCTGGCCACAAGCATTGCTTATGT<br>TTATTCTCTGGTCATCCTGGTGGTTGCTGTGGCTGAGAAGGCGGAGAGGAGCCCTGTGACA<br>TTCTTCGACACGCCCCCCATGCTCTTTGTGTTCATTGCCCTGGGCCGGTGGCTGGCAAAGGT<br>AACAGCAGCTTCAGGTTCAGAAAAGAGCTGCTCCTTCAGTAAACAAATCTCACTTCCTCTG<br>AACACCATGTTTAGAATTACTAATTATACACAGCATAGAGACAGACTTAAAGAAATAGGAAA<br>CCTCCATATAATTAAGGTGCTCTAGTCACTAATCTCCAAATTGGTCACTACTTCTGAAATCCC<br>AGCTAATCTGGTTAATTAT |
| 1-ins | CACTGTCCTTGTCTTTCAGCTCCTCGGTGGGTGGTACTTCTACGTTCAGGCCTACAAATCTC<br>TGAGACACAGGTCAGCCAACATGGACGTGCTCATCGTCCTGGCCACAAGCATTGCTTATGT<br>TTATTCTCTGGTCATCCTGGTGGTTGCTGTGGCTGAGAAGGCGGAGAGGAGCCCTGTGACA<br>TTCTTCGACACGCCCCCCATGCTCTTTGTGTTCATTGCCCTGGGCCGGTGGCTGGAACCACT<br>TGGCAAAGGTAACAGCAGCTTCAGGTTCAGAAAAGAGCTGCTCCTTCAGTAAACAAATCT<br>CACTTCCTCTGAACACCATGTTTAGAATTACTAATTATACACAGCATAGAGACAGACTTAAA<br>GAAATAGGAAACCTCCATATAATTAAGGTGCTCTAGTCACTAATCTCCAAATTGGTCACTAC<br>TTCTGAAATCCCAGCTAATCTGGTTAATTAT |
| 13-del | CACTGTCCTTGTCTTTCAGCTCCTCGGTGGGTGGTACTTCTACGTTCAGGCCTACAAATCTC<br>TGAGACACAGGTCAGCCAACATGGACGTGCTCATCGTCCTGGCCACAAGCATTGCTTATGT<br>TTATTCTCTGGTCATCCTGGTGGTTGCTGTGGCTGAGAAGGCGGAGAGGAGCCCTGTGACA<br>TTCTTCGACACGCCCCCCATGCTCTTTGTGTTCATTGCCCTGGGCCGGTGGCAAAGGTAAC<br>AGCAGCTTCAGGTTCAGAAAAGAGCTGCTCCTTCAGTAAACAAATCTCACTTCCTCTGAAC<br>ACCATGTTTAGAATTACTAATTATACACAGCATAGAGACAGACTTAAAGAAATAGGAAACCT<br>CCATATAATTAAGGTGCTCTAGTCACTAATCTCCAAATTGGTCACTACTTCTGAAATCCCAGC<br>TAATCTGGTTAATTAT |
| 2-del | CACTGTCCTTGTCTTTCAGCTCCTCGGTGGGTGGTACTTCTACGTTCAGGCCTACAAATCTC<br>TGAGACACAGGTCAGCCAACATGGACGTGCTCATCGTCCTGGCCACAAGCATTGCTTATGT<br>TTATTCTCTGGTCATCCTGGTGGTTGCTGTGGCTGAGAAGGCGGAGAGGAGCCCTGTGACA<br>TTCTTCGACACGCCCCCCATGCTCTTTGTGTTCATTGCCCTGGGCCGGTGGCTGGAACTTGG<br>CAAAGGTAACAGCAGCTTCAGGTTCAGAAAAGAGCTGCTCCTTCAGTAAACAAATCTCAC<br>TTCCTCTGAACACCATGTTTAGAATTACTAATTATACACAGCATAGAGACAGACTTAAAGAA<br>ATAGGAAACCTCCATATAATTAAGGTGCTCTAGTCACTAATCTCCAAATTGGTCACTACTTCT<br>GAAATCCCAGCTAATCTGGTTAATTAT |

Table S1. The gBlock synthetic ATP7B DNA sequences used for figure 1F.

| | WT | HDR | 9-del | 1-ins | 13-del | 2-del |
|---|---|---|---|---|---|---|
| 10 | 428.00<br>(418.54~437.46) | 44.55<br>(39.75~49.35) | 44.53<br>(38.93~50.12) | 53.38<br>(49.63~57.12) | 55.55<br>(48.93~62.17) | 45.73<br>(38.48~52.97) |
| 5 | 423.20<br>(416.84~429.56) | 21.44<br>(19.40~23.48) | 20.74<br>(18.44~23.04) | 25.48<br>(22.40~28.56) | 25.34<br>(22.17~28.51) | 21.52<br>(16.22~26.82) |
| 2.5 | 414.80<br>(400.66~428.94) | 10.48<br>(8.54~12.42) | 10.26<br>(7.46~13.06) | 12.69<br>(9.55~15.83) | 11.74<br>(8.55~14.92) | 10.66<br>(7.94~13.37) |
| 1.3 | 431.40<br>(414.11~448.69) | 5.41<br>(4.58~6.24) | 5.25<br>(4.33~6.17) | 6.83<br>(5.10~8.56) | 6.45<br>(4.30~8.60) | 5.08<br>(4.37~5.78) |
| 0.6 | 433.20<br>(419.39~447.01) | 2.52<br>(1.96~3.08) | 1.99<br>(1.36~2.62) | 2.45<br>(1.73~3.17) | 2.99<br>(1.83~4.16) | 2.64<br>(1.62~3.66) |
| 0.3 | 441.40<br>(438.05~444.75) | 1.16<br>(0.68~1.63) | 1.00<br>(0.27~1.73) | 1.53<br>(0.75~2.30) | 1.96<br>(1.21~2.72) | 1.22<br>(0.54~1.90) |
| 0.16 | 435.80<br>(419.72~451.88) | 0.80<br>(0.36~1.25) | 0.47<br>(0.18~0.75) | 0.91<br>(0.02~1.80) | 0.96<br>(0.32~1.60) | 0.63<br>(0.28~0.98) |
| 0.08 | 430.20<br>(415.01~445.39) | 0.41<br>(0.04~0.78) | 0.30<br>(-0.12~0.73) | 0.41<br>(0.01~0.82) | 0.79<br>(0.60~0.97) | 0.35<br>(0.15~0.55) |
| 0.04 | 440.00<br>(428.39~451.61) | 0.42<br>(0.15~0.68) | 0.14<br>(0.02~0.26) | 0.19<br>(-0.03~0.42) | 0.63<br>(0.41~0.86) | 0.04<br>(-0.02~0.10) |
| 0.02 | 433.80<br>(406.96~460.64) | 0.38<br>(0.22~0.53) | 0.07<br>(-0.02~0.16) | 0.10<br>(-0.01~0.21) | 0.28<br>(0.03~0.53) | 0.10<br>(0.02~0.19) |
| 0 | 434.60<br>(414.78~454.42) | 0.14<br>(0.06~0.22) | 0.03<br>(-0.05~0.11) | 0.02<br>(-0.03~0.06) | 0.45<br>(0.34~0.56) | 0.02<br>(-0.03~0.06) |

Table S2. Quantitative performance of the multi-color ddPCR assay assessed using serially diluted synthetic DNA templates.

Synthetic ATP7B DNA templates corresponding to the HDR allele and representative NHEJ alleles were spiked into 100 ng of wild-type genomic DNA at the indicated proportions and quantified by the multi-color ddPCR assay. Data are presented as mean measured concentrations (copies/mL) with 95% confidence intervals (95% CI) from replicate measurements. Values highlighted in red indicate dilution points for which the lower bound of the 95% CI overlapped with the upper bound of the 95% CI obtained from WT-only (0%) control samples. Limits of detection (LoD) were defined as the lowest input fraction whose 95% CI did not overlap with the corresponding WT-only control. Based on this criterion, the LoDs were determined to be 0.16% for HDR and 9-del, 0.3% for 1-ins and 13-del, and 0.08% for 2-del. n = 4 for the 10% dilution sample and n = 5 for all other dilution samples.

ATP7B

| | inDelphi | | ForeCasT | | NGS | |
|---|---|---|---|---|---|---|
| | Frequency | Rank | Frequency | Rank | Frequency | Rank |
| WT | – | – | – | – | 23.9 | 1 |
| HDR | – | – | – | – | 0.9 | 13 |
| 2-del | 22.1 | 1 | 34.5 | 1 | 17.3 | 2 |
| 13-del | 12 | 3 | 3.1 | 4 | 8.7 | 3 |
| 1-in | 12.9 | 2 | 3.5 | 2 | 8.4 | 4 |
| 9-del | 11.6 | 4 | 3.3 | 3 | 4.7 | 5 |
| Others | 41.4 | – | 55.6 | – | 36.1 | – |

RBM20

| | inDelphi | | ForeCasT | | NGS | |
|---|---|---|---|---|---|---|
| | Frequency | Rank | Frequency | Rank | Frequency | Rank |
| WT | – | – | – | – | 17.9 | 2 |
| HDR | – | – | – | – | 1.9 | 6 |
| 8-del_1 | 62.7 | 1 | 33.4 | 1 | 28.2 | 1 |
| 9-del | 1.9 | 8 | 5.3 | 2 | 14.3 | 3 |
| 21-del | 2.4 | 4 | 4.1 | 3 | 0.1 | 74 |
| 1-del | 1.3 | 10 | 2.6 | 4 | 5.6 | 4 |
| 1-in | 3.8 | 2 | 0.6 | 19 | 1.3 | 9 |
| 8-del_2 | 0.04 | 58 | 0.5 | 29 | 2.7 | 5 |
| Others | 27.86 | – | 53.5 | – | 29.3 | – |

Table S3. Comparison of predicted and experimentally observed genome editing outcomes.

The frequencies and rankings of NHEJ alleles predicted by inDelphi and FORECasT were compared with allele frequencies determined experimentally by next-generation sequencing (NGS) for ATP7B R778L and RBM20 R636S genome editing. WT and HDR allele frequencies are shown only for the NGS dataset. “Others” represents all remaining editing outcomes not included among the four highest-frequency predicted NHEJ alleles.

| Guide RNA | |
|---|---|
| ATP7B | GGGCCGGTGGCTGGAACACT |
| RBM20 | GGTCTCGTAGTCCGGTGAGC |
| APOE_gRNA-1 | GACATGGAGGACGTGTGCGG |
| APOE_gRNA-2 | TGGAGGACGTGTGCGGCCGC |
| Single strand oligo donor DNA (ssODN) | |
| ATP7B ssODN | CATGCTCTTTGTGTTCATTGCCCTGGGCCTGTGGCTGGAACACTTGGCAAAGGTAACAGC |
| RBM20 ssODN | ACAGATATGGCCCAGAAAGGCCGCGGTCTAGTAGTCCGGTGAGCCGGTCACTCTCCCCGA |

Table S4. Guide RNA and oligonucleotide sequences used for genome editing and base editing

ATP7B R778L 156 bp Assay Components

| Primers | Name | Sequence | | Final conc. |
|---|---|---|---|---|
| F | fw.2_ATP7B_R778L | TGCTTATGTTTATTCTCTGGTCATC | | 900 nM |
| R | rev.2_ATP7B_R778L | CCTGAAGCTGCTGTTACCTT | | 900 nM |
| Probes | Name | Sequence | Fluor-Quencher | Final conc. |
| Reference | Ref FAM probe | TGGTGGTTGCTGTGGCT | FAM-Zen | 250 nM |
| WT | WT HEX | CCGGTGGCTGGAACACT | HEX-Zen | 250 nM |
| HDR | HDR FAM | TGGGCCTGTGGCTG | FAM-Zen | 500 nM |
| NHEJ-1 | NHEJ 2-bp del ATTO590 | TGGCTGGAACTTGGCAAA | ATTO590-Iowa BlackRQ Sp | 250 nM |
| NHEJ-2 | NHEJ 13-bp del ROX | CGGTGGCAAAGGTAACAG | ROX-Iowa BlackRQ Sp | 250 nM |
| NHEJ-3 | NHEJ 1-bp ins Cy5.5 | CTGGAACCACTTGGCAAA | Cy5.5-Iowa BlackRQ Sp | 250 nM |
| NHEJ-4 | NHEJ 9-bp del Cy5 | GGTGGCTGGCAAAGGTAA | Cy5-TAO-Iowa BlackRQ Sp | 250 nM |

RBM20 R636S 150 bp Assay Components

| Primers | Name | Sequence | | Final conc. |
|---|---|---|---|---|
| F | fw.7_RBM20_R636 | CTGTGTGTGGGTGGGGT | | 900 nM |
| R | rev.7_RBM20_R636 | AGGAGGTGAAGCTGGGAG | | 900 nM |
| Probes | Name | Sequence | Fluor-Quencher | Final conc. |
| Reference | Ref FAM probe | TGGGAGGTGTGAAGATTCTAAATC | FAM-Zen | 250 nM |
| WT | WT HEX probe | CACTCGGCCAGTGAGA | HEX-Zen | 250 nM |
| HDR | WT DARK probe | CCGCGGTCTCGTAGTCC | FAM-Zen | 500 nM |
| HDR.dark | HDR FAM probe | CCGCGGTCTAGTAGTCC | None. Add 3' phosphate | 250 nM |
| NHEJ-1 | NHEJ 8-bp del_1 Cy5 probe | TCCGGTCACTCTCCCCGA | Cy5-TAO-Iowa BlackRQ Sp | 250 nM |
| NHEJ-2 | NHEJ 9-bp del ROX probe | CTCGTAGCCGGTCACTCTCC | ROX-Iowa BlackRQ Sp | 250 nM |
| NHEJ-3 | NHEJ 1-bp del Cy5.5 probe | TCTCGTAGTCCGGTAGCCG | Cy5.5-Iowa BlackRQ Sp | 250 nM |
| NHEJ-4 | NHEJ 8-bp del_2 ATTO590 probe | CTCGTAAGCCGGTCACTC | ATTO590-Iowa BlackRQ Sp | 250 nM |

APOE 252 bp Final Assay Components

| Primers | Name | Sequence | | Final conc. |
|---|---|---|---|---|
| F | APOE_ddPCR_Fw | TGGAGGAACAACTGACCC | | 900 nM |
| R | APOE_ddPCR_Rv2 | CAGCTCCTCGGTGCTCTG | | 900 nM |
| **Probes** | **Name** | **Sequence** | **Fluor-Quencher** | **Final conc.** |
| Reference | APOE Ref FAM probe | CTGTCCAAGGAGCTGCAG | FAM-Zen | 250 nM |
| APOE3 | APOE3 ATTO590 probe | CGGACATGGAGGACGTGTGCGG | ATTO590-Iowa BlackRQ Sp | 250 nM |
| APOE4 | APOE4 Cy5.5 probe | ACATGGAGGACGTGCGCGGCCG | Cy5.5-Iowa BlackRQ Sp | 500 nM |
| Bystander1 | Bystander1 HEX probe | ACATGGAGGACGCGCGCGGCCG | HEX-Zen-Iowa BlackFQ | 250 nM |
| Bystander2 | Bystander2 Cy5 probe | ACATGGAGGACGCGTGCGGCCG | Cy5-TAO-Iowa BlackRQ Sp | 250 nM |

Table S5. Primer and probe sequences used for multi-color ddPCR assays.

Sequences, fluorophore-quencher combinations, and final reaction concentrations of primers and probes used in the ATP7B R778L, RBM20 R636S, and APOE multi-color ddPCR assays. All probes were synthesized by Integrated DNA Technologies. Final concentrations correspond to those used in ddPCR.
